# Macrolide resistance through uL4 and uL22 ribosomal mutations in *Pseudomonas aeruginosa*

**DOI:** 10.1101/2024.04.11.588999

**Authors:** Lise Goltermann, Pablo Laborda, Oihane Irazoqui, Ivan Pogrebnyakov, Søren Molin, Helle Krogh Johansen, Ruggero La Rosa

## Abstract

Macrolides are widely used antibiotics for the treatment of bacterial airway infections. Due to its elevated minimum inhibitory concentration in standardized culture media, *Pseudomonas aeruginosa* is considered intrinsically resistant and, therefore, antibiotic susceptibility testing against macrolides is not performed. Nevertheless, due to macrolides’ immunomodulatory effect and suppression of virulence factors, they are used for the treatment of persistent *P. aeruginosa* infections. Here, we demonstrate that macrolides are, instead, effective antibiotics against *P. aeruginosa* airway infections in an air-liquid interface (ALI) infection model system resembling the human airways. Importantly, macrolide treatment in both people with cystic fibrosis and primary ciliary dyskinesia patients leads to the accumulation of uL4 and uL22 ribosomal protein mutations in *P. aeruginosa* which causes antibiotic resistance. Consequently, higher concentrations of antibiotics are needed to modulate the macrolide-dependent suppression of virulence. Surprisingly, even in the absence of antibiotics, these mutations also lead to a collateral reduction in growth rate, virulence and pathogenicity in airway ALI infections which are pivotal for the establishment of a persistent infection. Altogether, these results lend further support to the consideration of macrolides as *de facto* antibiotics against *P. aeruginosa* and the need for resistance monitoring upon prolonged macrolide treatment.

## Introduction

Macrolides are widely used antibiotics for the treatment of several bacterial airway infections. They inhibit protein synthesis by binding the antibiotic binding site within the nascent polypeptide exit tunnel (NPET) of the bacterial ribosome. This modifies the translational capacity of the cell and blocks its growth ^1–3^. Together with their bacteriostatic effect, in several bacterial species, macrolide antibiotics reduce the expression of virulence factors by a yet unclear mechanism ^4–6^. Moreover, macrolides provide immunomodulatory effects on the host which benefit a large group of patients with inflammatory respiratory diseases ^7–9^. In people with cystic fibrosis (pwCF), primary ciliary dyskinesia (PCD) and chronic obstructive pulmonary disease (COPD), macrolides are prescribed as continuous low-dose treatment to reduce the risk of exacerbations and limit the host immune response against bacterial pathogens ^10,11^. However, whether such prolonged courses of antibiotics are contributing to the development of antibiotic resistance is not always determined. For instance, according to the European Committee on Antimicrobial Susceptibility Testing (EUCAST), *Pseudomonas aeruginosa* is classified as intrinsically resistant towards macrolide antibiotics and, therefore, not susceptible ^12,13^. Consequently, antibiotic susceptibility testing (AST) of clinical *P. aeruginosa* isolates is not carried out for this drug class although macrolide treatment is routinely prescribed in pwCF, PCD and COPD patients chronically infected with *P. aeruginosa* ^13^. Therefore, no consideration has been given to the possibility that macrolides are, in fact, effective antibiotics against *P. aeruginosa* and to the consequent resistance development in clinical strains ^14^.

Recent implementations of *in vivo*-like model of airway epithelium in an Air-Liquid Interphase (ALI) have disclosed new mechanisms of antibiotic resistance that could not be identified by classical Minimum Inhibitory Concentration (MIC) evaluations ^15^. Indeed, the localization of the bacteria within the epithelium, the ability of the antibiotics to cross the cell layer and the availability of nutrients can modify the bacterial susceptibility. This is not evident from standardized MIC assays, but it requires taking into account the complexity of the host environment including the relevant interactions between the host and the pathogen. Recently, it has been discovered that alterations to the antibiotic susceptibility medium for *P. aeruginosa* testing reduce the MIC of macrolides significantly ^14,16^. Diluted LB medium or cell culture media (RPMI medium) might mimic the metabolic demands of infecting bacteria *in vivo* better than other standardized AST media such as Müeller-Hinton medium. This has allowed for the identification of strains with decreased macrolide susceptibility, paving the way for more thorough investigations of once unattainable mechanisms of resistance ^14^. This strongly suggests that macrolides are, in fact, effective antibiotics against *P. aeruginosa*. However, a systematic analysis of mechanisms of macrolide resistance is still lacking. Furthermore, limited information is available on their activity *in vitro* and/or *in vivo* during an infection and on the consequences of prolonged treatments.

Previously, we identified and characterized clinical *P. aeruginosa* isolates from pwCF which were significantly less susceptible to erythromycin and azithromycin than their most closely related ancestors ^14^. In addition, macrolide-induced effects on redox sensitivity and motility were also reduced. This was caused by mutations in the uL4 ribosomal protein which together with the uL22 ribosomal protein and other ribosomal proteins, form the NPET of the ribosome close to the antibiotic binding site ^17^. Similar mutations have been identified in other bacterial species susceptible to macrolide antibiotics such as *Escherichia coli*, *Streptococcus pneumoniae*, *Legionella pneumophila* and *Neisseria gonorrhoeae* ^18,19^. Moreover, mutations in the 23S rRNA have also been identified as drivers of resistance to macrolides in several clinical strains ^20,21^. However, thorough investigations of mechanisms of resistance have not been carried out in *P. aeruginosa* because of its perceived intrinsic macrolide resistance. Moreover, it is often challenging to evaluate the effect of ribosomal mutations on antibiotic susceptibility in clinical isolates. This is because clinical isolates harbour a high genomic complexity due to a large number of mutations with epistatic interactions, which drastically modify the bacterial phenotype. Therefore, whether *P. aeruginosa* is intrinsically susceptible to macrolides, whether mutations in the uL4 and uL22 ribosomal proteins provide increased antibiotic resistance and if such mutations lead to other secondary effects remains unclear. Ribosomal mutations can have unforeseen consequences due to the key function of the translation machinery ^1^. For example, they can change the translational landscape and induce or abolish ribosomal pausing at certain sequence motifs providing changes in gene expression ^22,23^. Moreover, whether macrolide-dependent virulence modulation is dependent on the specific historical contingency of the clinical isolates or if such effect is generalizable, is largely unclear.

We hypothesize that macrolide antibiotics are in fact effective against *P. aeruginosa*^16^. This in turn warrants investigations of possible resistance mechanisms and their effect on growth, fitness, phenotype and virulence in the presence or absence of antibiotics. Furthermore, we hypothesize that macrolide treatment in pwCF and PCD patients exerts a selective pressure for the generation of an increasing number of macrolide-resistant *P. aeruginosa* strains. Unfortunately, these strains might go unnoticed in the laboratory as a result of the wrong assumption that *P. aeruginosa* has an inherent resistance to macrolides. Here, we use a reverse engineering approach by taking a selection of previously identified uL4 and uL22 mutations and moving each of them into the same genetic background of a reference strain. This isolates the effects of the ribosomal mutations from the tens or hundreds of other mutations found in the clinical isolates. Moreover, it allows for a direct comparison of susceptibility, growth rate, proteome allocation and virulence between the mutants. To assess the biological consequences of uL4 and uL22 ribosomal mutations during an infection, we performed infection assays in an ALI model system of human airway bronchial epithelium. To further establish the clinical relevance of ribosomal protein mutations causing macrolide resistance, we analysed 663 newly sequenced *P. aeruginosa* isolates from pwCF as well as patients with PCD to identify any accumulated uL4 and uL22 protein mutations causing macrolide resistance.

## Results

### uL4 and uL22 mutations confer antibiotic resistance and reduced virulence modulation

Several uL4 and uL22 mutations conferring resistance to macrolide antibiotics were previously identified in clinical strains of *P. aeruginosa* ^14^. Since clinical isolates have complex historical contingency including a multitude of mutations interacting epistatically, we aimed to characterize the effect of uL4 and uL22 mutations upon introduction into the same genetic background, in this case, that of the laboratory strain PAO1.

By employing a CRISPR-Cas system ^24^, we successfully introduced three uL4 and four uL22 mutations into the *P. aeruginosa* PAO1 reference strain genome which were further verified by WGS (Supplementary Data 1). The three uL4 mutations, 1′KPW (1′57-KPW-59), 1′RA (1′68-RA-69) and +G (+G65), were initially identified in the clinical collection of *P. aeruginosa* isolates from the CF clinic at Rigshospitalet, Copenhagen, Denmark. The four uL22 mutations 1′IV (1′96-IV-97), 1′KRI (1′83-KRI-85), 1′VKRS (1′97-VKRS-100), and +GK (+89-GK-90) were, instead, found among sequenced *P. aeruginosa* strains from different infection scenarios and collections ^14^.

For all uL4 and uL22 mutants, the susceptibility to the macrolides erythromycin and azithromycin was significantly reduced compared to the parental PAO1 strain (Fig. 1A). Erythromycin’s MIC increased from 128 μg/ml for the wild-type PAO1 to 512-1024 μg/ml for the uL4 and uL22 mutants. Resistance to azithromycin increased dramatically, with MIC values going from 16-32 μg/ml (wild-type PAO1) to 128 μg/ml (uL4 mutants) and 512 μg /ml (uL22 mutants). A similar result was obtained when analyzing the viability of bacteria after the MIC assay (Post-MIC effect). Indeed, the concentration of antibiotic needed to fully inhibit the growth was significantly higher for the uL4 and uL22 mutants compared to the PAO1 wild-type (Fig. 1B).

**Figure 1.**
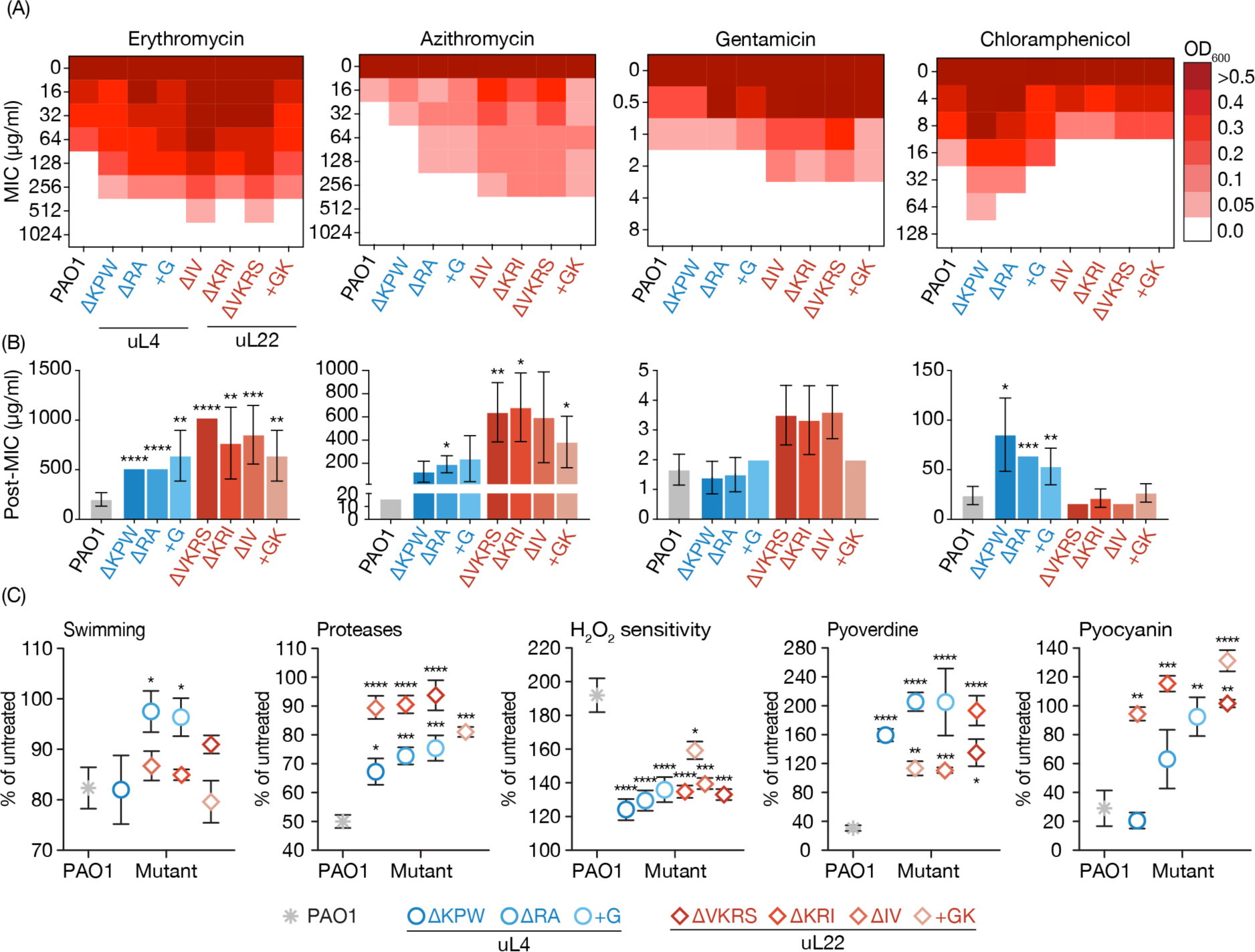
Antibiotic susceptibility testing and macrolide-dependent phenotype modulation for uL4 and uL22 *P. aeruginosa* mutant strains. (A) The minimum inhibitory concentration (MIC) was determined by endpoint optical density (OD_630_) after 24h incubation in a MIC assay in 50% LB with increasing concentration of antibiotic. (B) Post-MIC effect was determined as the minimum concentration needed to prevent re-growth when spotted onto LB-agar after 24h MIC incubation. MIC and Post-MIC were compared between strains for erythromycin, azithromycin, gentamicin and chloramphenicol. (C) The increase or decrease in swim diameter, protease secretion, redox (H_2_O_2_) sensitivity, pyoverdine and pyocyanin production was computed as a percentage (%) between untreated and azithromycin treated (2% of the mutant MIC) cultures. The data represent the mean ± SD of 3-6 biological replicates. Statistical significance relative to PAO1 wild-type was computed by One-Way ANOVA followed by the Dunnett multiple comparisons test where *P < 0.05, **P < 0.01, ***P < 0.005, ****P < 0.001.

Next, we tested collateral sensitivity/cross-resistance against gentamicin and chloramphenicol, which target other sites on the ribosome. The uL22 mutants resulted slight increase in gentamicin MIC, which was most likely caused by the reduced growth rate of some of the mutant strains ^25^ (Fig. 1A and B). In contrast, each of the four uL4 mutants was significantly less susceptible to chloramphenicol when compared with the wild-type PAO1 and the uL22 mutant strains (Fig. 1A and B). This could suggest that the uL4 deletions cause a certain degree of rearrangement of the ribosome’s structure, the effect of which extends to the chloramphenicol binding site ^26^. Instead, a panel of non-ribosome targeting antibiotics did not show any significant differences in the MIC for any strains, compared to the PAO1 wild-type (Table S1).

Together with its bacteriostatic effect, macrolides exert an anti-virulence effect suppressing quorum sensing dependent virulence ^4,6,27^. Therefore, we quantified various parameters such as swim diameter, protease secretion, redox sensitivity, pyoverdine and pyocyanin production in the absence or presence of sub-MIC concentrations of azithromycin for all strains (Fig. 1C). As expected, most of the tested phenotypes were significantly less affected by the presence of the antibiotic in the mutant strains when compared to PAO1 (Fig. 1C).

These results confirm that each of the selected mutations did in fact confer increased macrolide resistance. Furthermore, changes in antibiotic susceptibility only apply to ribosome-targeting compounds, ruling out collateral sensitivity or cross-resistance toward other drug classes. Lastly, azithromycin exerts reduced suppression of virulence traits as a consequence of the reduced susceptibility of uL4 and uL22 mutant strains.

### Changes in phenotype and growth physiology in uL4 and uL22 mutant strains

Ribosomal mutations often cause fitness costs due to impaired ribosome assembly or suboptimal protein translation rates. This is often shown as reduced growth abilities and altered cell physiology ^28^. While under normal growth conditions, growth defects might not be significant, possible defects in ribosome assembly are frequently worsened at lower temperatures ^29^. To account for this, growth rates were determined and compared between 37°C and 30°C as well as with increasing concentrations of erythromycin.

At 37°C and in the absence of antibiotics, the ΔRA (9.8%), ΔVKRS (46.1%), ΔKRI (31.7%) and 1′IV (41.5%) showed statistically significant reduced growth rates compared to the wild-type PAO1 (Fig. 2A). As expected, the growth rate of PAO1 decreased rapidly with increasing erythromycin concentration at 37°C as well as 30°C. Similar profiles were observed with the +G (uL4) and +GK (uL22) insertion mutants, suggesting a comparable ribosome functionality (Fig. 2A). Instead, the ΔKPW (48%) and ΔRA (31%) deletion mutants, showed a higher reduction of growth rate from 37°C to 30°C compared to the reduction seen in PAO1 (26%) (Fig. 2A). This might suggest a possible ribosome assembly/translation defect which is evidenced when reducing the temperature to 30°C. In contrast, all the uL22 deletion mutants showed a reduced growth rate (range 31.7% - 46.1%) compared to PAO1 but were, however, generally less affected by erythromycin treatment or temperature reduction (Fig. 2A). This effect was most prominent at lower temperature where the addition of antibiotic in the culture medium did not alter the strains’ growth rate (Fig. 2A).

**Figure 2.**
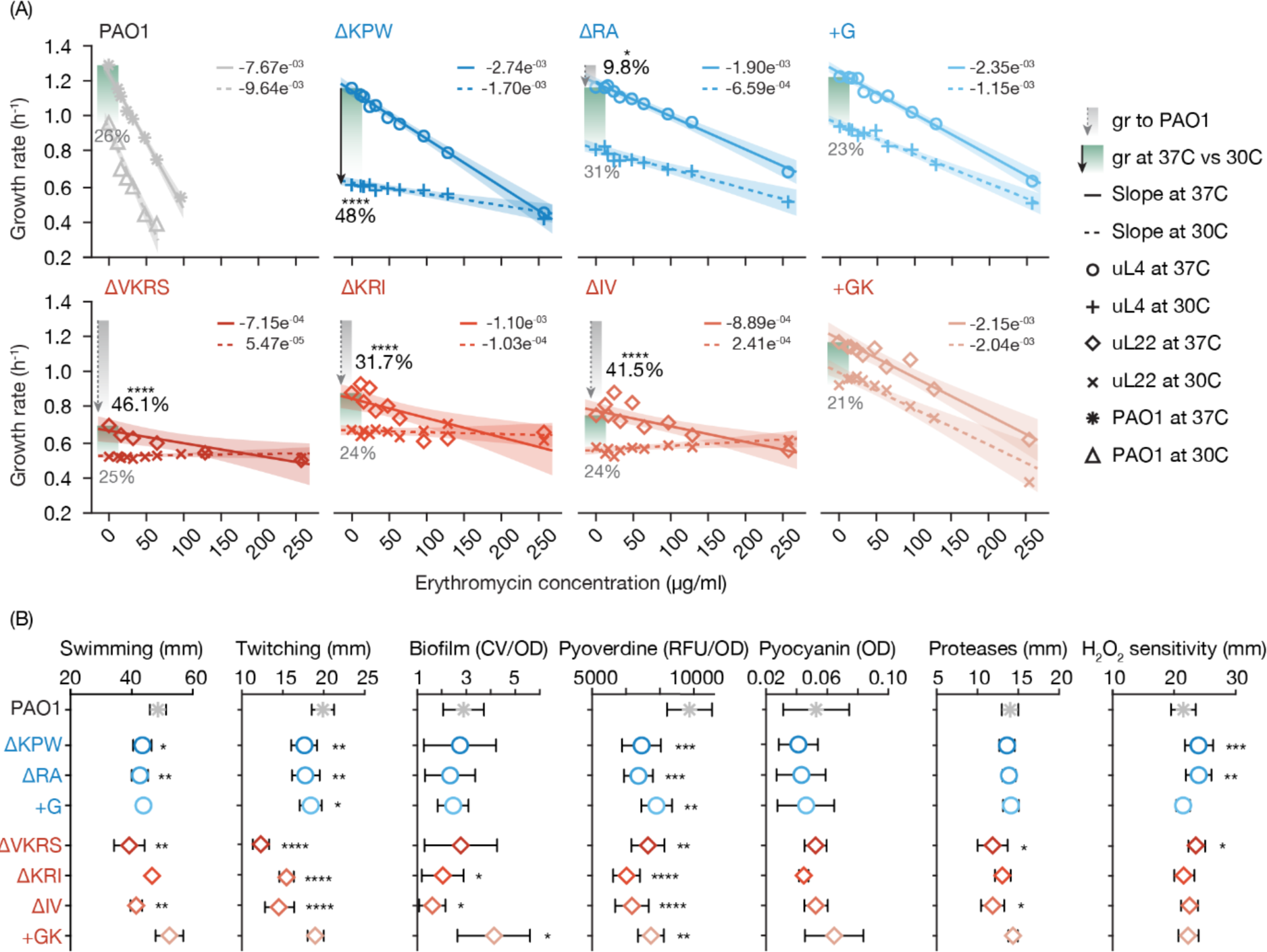
Phenotype profiling of *P. aeruginosa* uL4 and uL22 mutant strains. (A) Maximum growth rates were calculated at 37°C and 30°C with increasing concentrations of erythromycin. Grey bars indicate growth reduction relative to PAO1 at 37°C. Green bars show growth reduction between 37°C and 30°C. (B) Swimming and twitching diameter, biofilm production, pyoverdine and pyocyanin production, protease secretion and redox (H_2_O_2_) sensitivity were tested for all ribosomal mutants and compared with the parental PAO1 wild-type strain. The data represent the mean ± SD of 3-6 biological replicates. Statistical significance relative to PAO1 wild-type and for the 37°C vs 30°C comparison was computed by One-Way ANOVA followed by Dunnett multiple comparisons test where *P < 0.05, **P < 0.01, ***P < 0.005, ****P < 0.001.

In addition to altering the growth physiology of the cell, ribosomal mutations may also alter other phenotypic traits independent of antibiotic susceptibility. Motility (swimming and twitching), biofilm formation, secondary metabolites secretion (pyoverdine and pyocyanin), protease secretion and redox sensitivity were, therefore, investigated for all uL4 and uL22 mutants in the absence of antibiotics. No substantial differences between the mutant strains and the PAO1 wild-type strain were observed for any of the tested phenotypes (Fig. 2B). Indeed, some small variations may be attributable to differences in growth rate as for example the reduction in swimming and twitching motility for the uL4 and uL22 deletion strains. Only pyoverdine production seems to be consistently reduced for all ribosomal mutants when compared to the PAO1 wild-type strain (Fig. 2B).

These results indicate that the deletions in the uL4 and uL22 ribosomal proteins result in fitness-associated costs such as cold sensitivity, assembly/translation defects, and slight changes in bacterial phenotype. Insertion mutants (+G and +GK) have, instead, no apparent fitness cost and a wild-type like growth physiology and phenotype.

### Proteome analysis reveals 3 distinct groups of functionally distinct mutations

To determine whether ribosomal mutations affected ribosome functionality and cellular expression, we compared the proteome composition of all mutant strains to that of the PAO1 wild-type.

Both principal component analysis (PCA) and hierarchical clustering analysis (HCA) separated the proteomes into three groups based on the type of mutation: i) uL4 deletions, ii) uL22 deletions and iii) uL4 and uL22 insertions mutants together with PAO1 wild-type (Fig. 3A and B). Similarly, among the three groups, uL22 deletion strains showed the highest number of differentially expressed proteins, followed by the uL4 deletion mutant and finally by the uL4 and uL22 insertion mutants (Fig. 3C). This grouping is also noticeable in the growth characteristics and physical traits of the strains. The uL22 deletion mutants display a reduced growth rate, while the uL4 deletions exhibit cold-sensitivity. The insertion mutants, on the other hand, show similar growth rates as the wild-type PAO1 (Fig. 2A).

**Figure 3.**
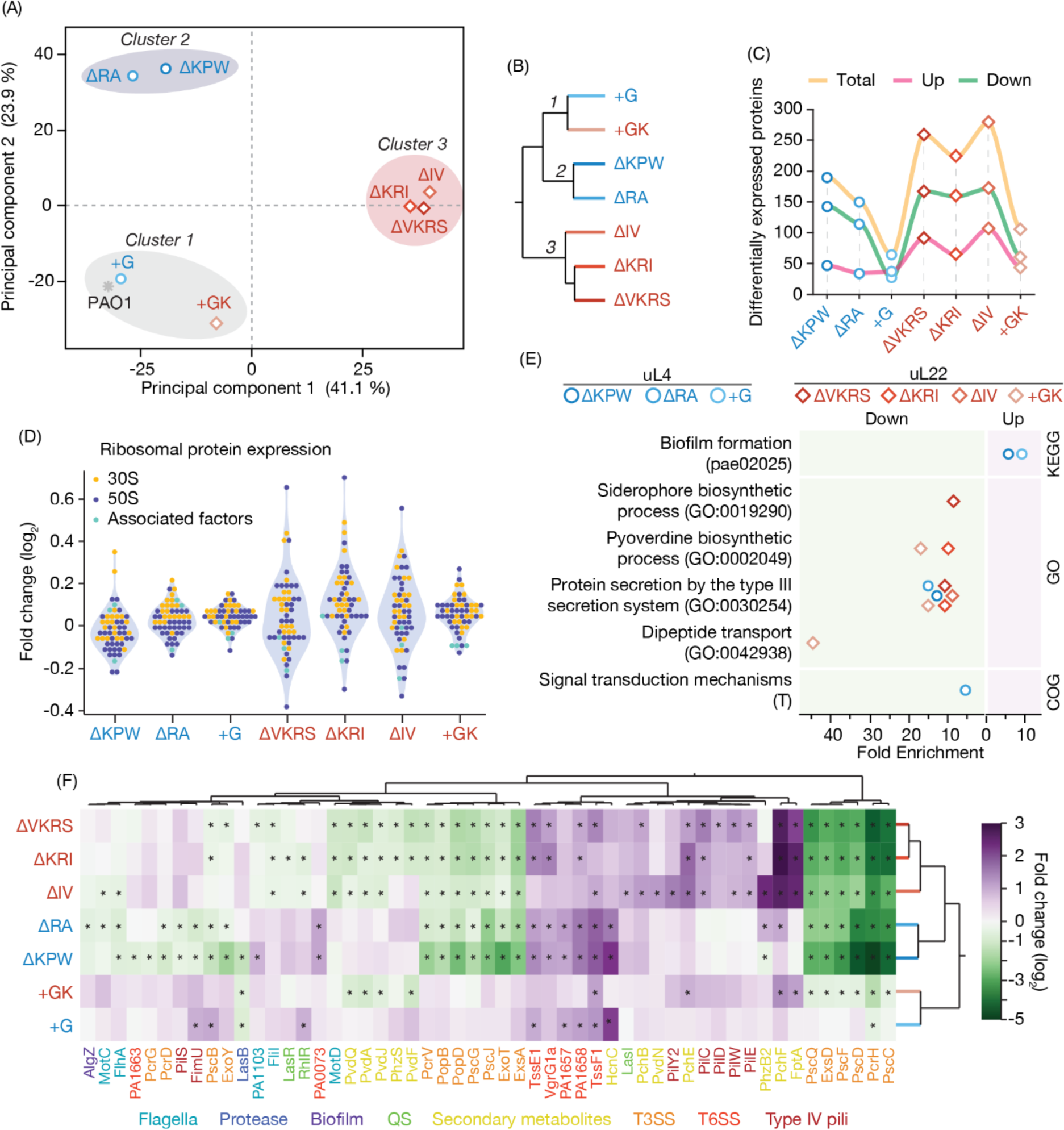
Proteome composition analysis of the *P. aeruginosa* wild-type PAO1 and the uL4 and uL22 mutant strains. (A) Principal Component Analysis of the proteomes based on the normalized expression of 2595 proteins identified. (B) Hierarchical Clustering Analysis based on the 473 differentially expressed proteins in the comparisons between mutant strains *vs* PAO1 wild-type. (C) Number of differentially expressed proteins for each comparison. The orange line indicates the total number of differentially expressed proteins, the purple line the number of up-regulated proteins and the green line the number of down-regulated proteins. (D) Expression profile of ribosomal proteins. The graph shows the log_2_ fold-change of the comparison mutants *vs* PAO1 wild-type. (E) Enrichment analysis of the strain-specific differentially expressed proteins (up- and down-regulated) for the COG, GO and KEGG terms. (F) Normalized expression profile (log_2_ fold-change) of the virulence determinant according to the Virulence Factor Database, Victors Virulence Factors and the Pseudomonas Genome Database (PseudoCAP). The asterisk indicates statistical significance in the comparison of mutants *vs* PAO1 wild-type.

Analyzing specifically ribosomal proteins and their associated factors showed no significant change in their abundance in either mutant (fold-change > ± 0.6; *P* value < 0.05). This suggests that ribosomal proteins do not require major compensatory expression changes to accommodate the uL4 and uL22 mutations. However, there is a higher variation in the expression of single ribosomal proteins and associated factors in the uL22 deletion mutants compared to the other mutants (Fig. 3D). This may suggest that ribosome reorganization might be necessary for its proper assembly, which is well in line with the observed reduced growth rate in these mutants.

Interestingly, analyzing the clusters of orthologous groups (COG) categories of differentially expressed proteins to identify underlining changes in proteome composition, showed that several proteins belonged to the categories of amino acid transport and metabolism and defense mechanisms (Supplementary Data 1). Furthermore, categories of differentially expressed proteins statistically enriched within the Gene Ontology (GO) classification included: i) the down-regulation of the protein secretion by the type III secretion system (T3SS) (6 of 7 strains); ii) the down-regulation of the proteins involved in pyoverdine biosynthetic processes (2 of 7 strains); iii) the up-regulation of proteins involved in biofilm formation (2 of 7 strains) (Fig. 3E). While changes in metabolic functions can be explained by differences in growth rate and growth physiology between strains, changes in virulence and defense determinants suggest a secondary effect of ribosomal mutations beyond antibiotic resistance. Indeed, when examining more broadly proteins involved in virulence and pathogenicity, we found a reduction in the expression of T3SS and flagella in the uL4 and uL22 deletion mutant strains. Conversely, we observed an increase in the expression of type 6 secretion system (T6SS), type IV pili, and secondary metabolites (Fig. 3F). In *P. aeruginosa*, the T3SS and the T6SS have a key role during the early colonization phase since they modify the host cell machinery influencing both invasion, growth, and host immune response ^30^. In the insertion mutant +G and +GK, instead, virulence proteins were expressed at a similar level to PAO1 indicating a limited effect of these mutations in pathogenicity. However, it should be noted that the growth conditions for the proteomic analysis did not promote virulence determinants specifically. This implies that the effect of ribosomal protein mutations on virulence suppression can differ *in vivo* during an infection, affecting a broader or narrower range of factors.

These results indicate that mutant strains change their proteome composition according to the specific mutation (uL4 deletions *vs* uL22 deletions *vs* uL4 and uL22 insertions). This effect goes beyond that of the ribosome functionality and antibiotic resistance, likely involving virulence determinants and pathogenicity.

### Azithromycin exerts an antibiotic effect in Air-Liquid Interface infection assays

AST, phenotypic characterization, and proteomic analyses all indicated that the uL4 and uL22 mutant strains have reduced susceptibility to macrolide antibiotics and modified expression of virulence determinants in laboratory conditions. To confirm these results in *in vivo*-like conditions, we performed infection assays in an ALI infection model system of pseudostratified airway epithelium. This system allows for the evaluation of both the host response to an infection and the virulence of the infecting pathogens. We selected the 1′RA and the +G strains (uL4 mutants) along with PAO1 wild-type since in laboratory conditions they have similar growth rates but different proteomic profiles. This allows for the direct comparison of the bacterial virulence during the infection, and it avoids misinterpretations caused by secondary effects such as the reduced growth rate. As a negative control of the infection, we used a Δ*pscC* mutant strain, which has an impaired T3SS and is susceptible to macrolides.

First, we tested the antibiotic efficacy of azithromycin in the ALI model when infected with either *P. aeruginosa* strain. Bacteria were allowed to establish an infection for 7h (pre-incubation) after which azithromycin treatment was applied for 14 h (Azi treatment). We measured the transepithelial electrical resistance (TEER), which quantifies the integrity and permeability of the epithelial layer, LDH release, which quantifies the epithelium cellular damage, and the bacterial count which quantifies the growth and penetration of the bacteria through the epithelium to the basolateral side of the ALI transwells. All measured parameters confirmed that while comparable values were shown between strains during the pre-incubation phase in absence of antibiotics, the ΔRA and the +G strains showed a significant damage to the epithelial layer and sustained bacterial growth after azithromycin treatment (Fig. 4A-C). Instead, the wild-type PAO1 and the Δ*pscC* mutant strain showed only limited growth and cellular damage due to the inhibitory effect of azithromycin on the bacterial growth (Fig. 4A-C). These results are corroborated by the confocal microscopy images which show substantial growth for the ΔRA and the +G strain infections and growth inhibition for the PAO1 and Δ*pscC* strains (Fig. 4D).

**Figure 4.**
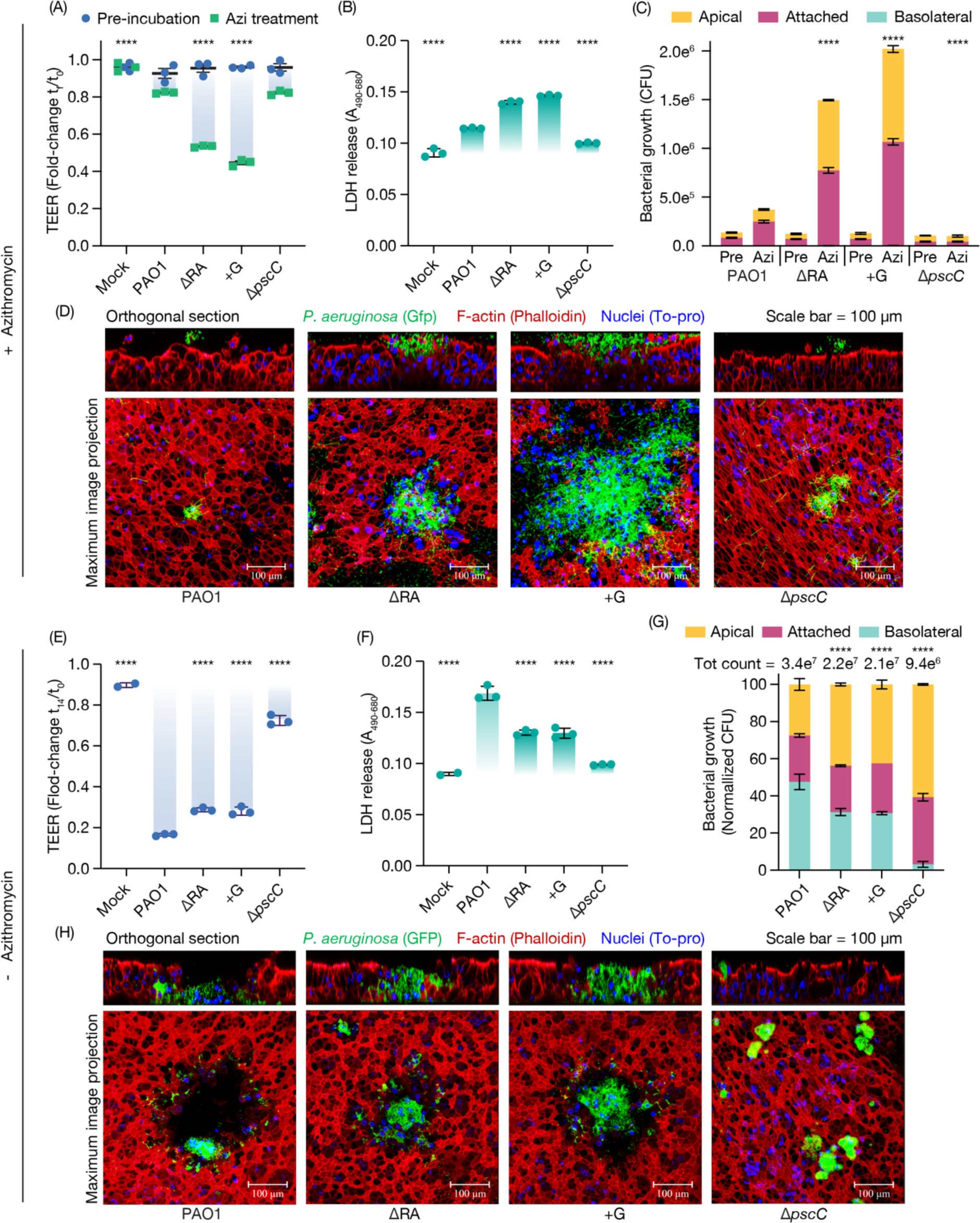
Antibiotic susceptibility and bacterial virulence in Air-Liquid Interface infection model. (A and E) Transepithelial Electrical Resistance (TEER) upon infection with strains PAO1, 1′RA, +G and 1′*pscC* in (A) presence and (E) absence of azithromycin. The data show the mean ± SD of the fold-change of the final TEER (t_f_) relative to time 0. The pre-incubation phase lasted 7 h following a 14 h azithromycin treatment. For all experiments, three biological replicates were used, and azithromycin concentration was set to 512 μg/ml. (B and F) LDH release by damaged epithelial cells upon infection in (B) the presence and (F) absence of azithromycin. (C) Bacterial growth in the apical side (yellow), attached to the epithelium (pink) and in the basolateral side (green) in (C) presence and (G) absence of azithromycin. In (G), bacterial growth is shown as % of the total count presented on top of the graph. (D and H) Confocal microscopy images of ALI transwells following infection with *P. aeruginosa* strains in green (GFP), epithelium in red (Phalloidin) and nuclei in blue (To-pro) in (D) presence and (H) absence of azithromycin. In all cases, statistical significance relative to PAO1 wild-type was computed by One-Way ANOVA followed by Dunnett multiple comparisons test where *P < 0.05, **P < 0.01, ***P < 0.005, ****P < 0.001. In (A and C) statistical significance was computed by Two-Way ANOVA to compare differences between pre-incubation and azithromycin treatment.

Next, we tested the virulence of the uL4 mutant strains in the absence of any antibiotic to evaluate the role of reduced virulence determinant expression during an infection. Interestingly, we found that both 1′RA and +G strains caused reduced cellular damage relative to PAO1. This is in line with the 1′*pscC* strain which has a significantly reduced penetration into the epithelial layer due to the absence of a functional T3SS (Fig. 4E and F). Similarly, the bacterial growth is reduced in the 1′RA and the +G and 1′*pscC* mutant strains (total count) since accessibility to the nutritional resources present in the basolateral compartment is limited because of the reduced penetration (Fig. 4G). It is worth noting that the bacterial growth of the 1′RA and the +G and 1′*pscC* is comparable to that of PAO1 in laboratory conditions (Fig. 2A), confirming that the reduced cellular damage and penetration, even if subtle, is driven by the reduced expression of virulence traits in the uL4 mutant strains as indicated by the proteomic analysis (Fig. 3F). This is also evidenced by the distinct distribution of the bacteria preferentially located on the apical side and attached to the epithelium for the 1′RA, +G and 1′*pscC* while largely present in the basolateral side in the wild-type PAO1 (Fig. 4F). As expected, instead, the PAO1 wild-type infections led to larger perforations and consequent degradation of the airway epithelium than that of the uL4 and 1′*pscC* mutant strains (Fig 4H). Importantly, we speculate that in ALI conditions the +G mutant has a comparable proteomic and virulence profile to that of the 1′RA strain since no differences in infection pattern were shown between the strains.

Altogether these results suggest a dual effect of the uL4 ribosomal mutations leading to macrolide reduced bacteriostatic effect and reduced penetrations and cellular damage which is an advantage for the maintenance of a long-term infection.

### Persistence of uL4 and uL22 mutations in pwCF and PCD

Due to the wrong assumption of *P. aeruginosa*’s intrinsic macrolide resistance, limited consideration has been given to the possibility of resistance development in clinical strains. Given that macrolides are instead effective antibiotics against *P. aeruginosa*, we expect that macrolide treatment exerts a selective pressure for the generation of macrolide-resistant strains. To confirm this hypothesis, we analysed 663 newly sequenced *P. aeruginosa* isolates from pwCF and patients with PCD to identify potential uL4 and uL22 protein mutations causing macrolide resistance. Importantly, in addition to pwCF, guidelines now also recommend long courses of azithromycin treatment in PCD patients ^10^.

From 431 isolates collected from 67 pwCF and 232 isolates collected from 64 patients with PCD treated with azithromycin, we identified five different uL4 mutations and four different uL22 mutations. uL4 mutations were present in 20 isolates from 7 pwCF and 1 PCD patient. uL22 mutations were found in 15 isolates from 2 pwCF and 1 PCD patient (Fig. 5A and Table S2). The mutations consist of 3 deletions, 3 insertions, and 3 substitutions. Most of the uL4 mutations are also present in other strain collections ^14^ (Table S2). The +KG insertion has not been identified before, but it is found in both a pwCF as well as a person with PCD. The four uL22 mutations are unique, and they map to the same area in the β hairpin loop of uL22 as previously described ^14^ (Fig. 5A and Table S2). The mutations are distributed over a range of different clone types, defined as lineages which differ by > 10,000 SNPs (Fig. 5A and Table S2). This suggests that such mutations can be accommodated in different genetic backgrounds. While some mutations were only identified in a single isolate, most mutated isolates persisted between 2 (G>A) and 8 years (+RK) in each patient (Fig. 5A and Table S2) underlining the fact that these mutants are highly adaptive to the CF and PCD airways and can persist long term.

**Figure 5.**
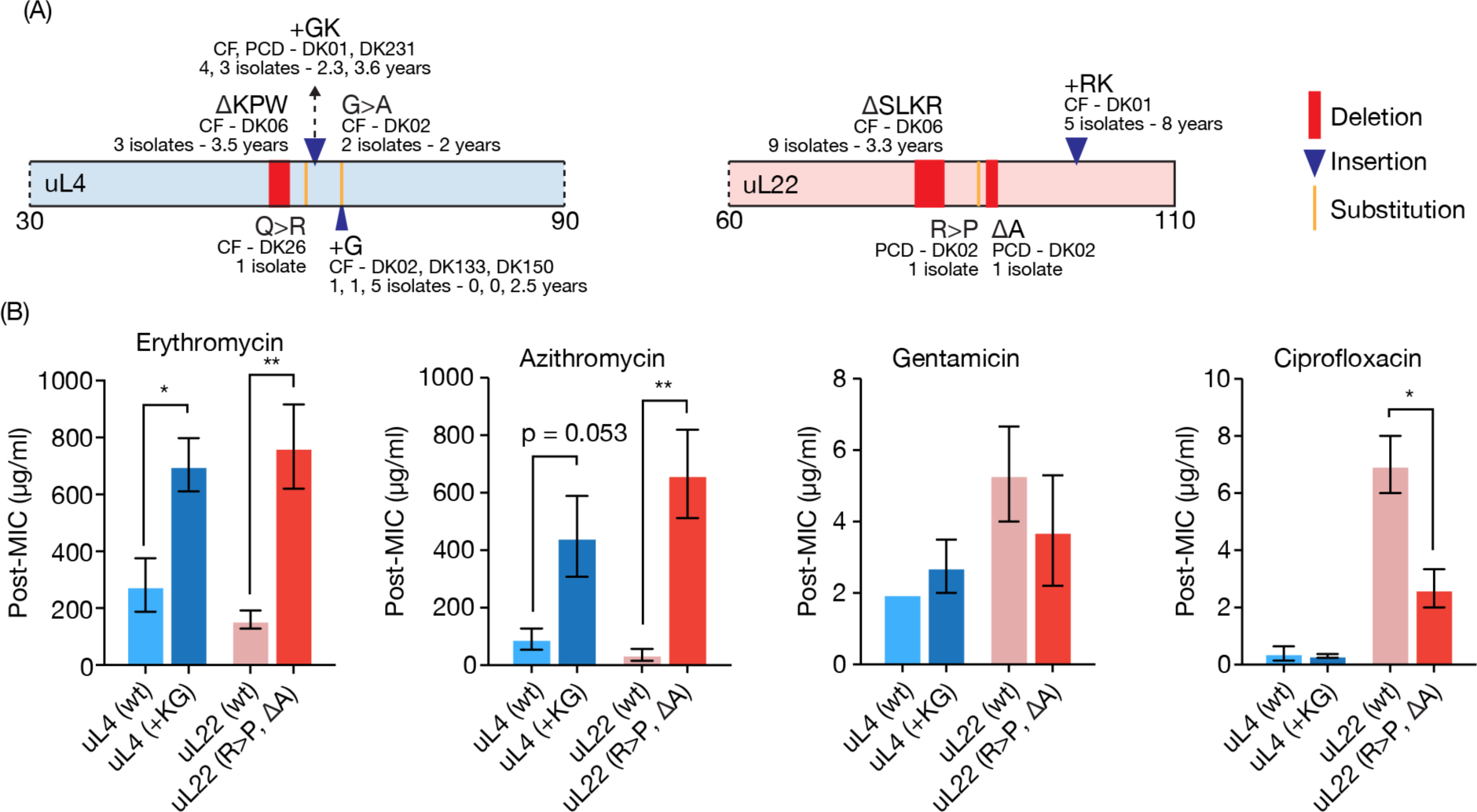
Identification of uL4 and uL22 mutations in *P. aeruginosa* isolates from pwCF and PCD. (A) Schematic representation of the distribution of mutations (red rectangle deletions, blue triangle insertions and yellow bar substitutions) in the uL4 and uL22 proteins. The specific mutation, origin of the isolate, the clone type, the number of isolates and the persistence are indicated below each mutation. (B) The minimum inhibitory concentration (MIC) for azithromycin, erythromycin, gentamicin and ciprofloxacin was determined by growth after re-plating (Post-MIC) after 24h incubation in a MIC assay in 50% LB with increasing concentration of antibiotic. For each mutant strain, the MIC was compared to an ancestor without the mutation from the same patient. The data represent the mean ± SD of 3-6 biological replicates. Statistical significance relative to each ancestor was computed by Student’s t-test where *P < 0.05, **P < 0.01, ***P < 0.005, ****P < 0.001.

AST was conducted on two strains from PCD patients containing the mutations +KG (uL4) and R>P, 1′A (uL22). The results were compared to the respective ancestor strain from the same clone type, which was isolated from the same patient before the occurrence of the ribosomal mutation. The testing showed that each mutant isolate displayed a 2-4 fold increase in MIC for erythromycin and a 4-8 fold increase for azithromycin (Fig. 5B). The MICs for gentamicin and ciprofloxacin were, instead, not increased relative to the respective ancestor strain, thus confirming the specific effect of the mutations on macrolide susceptibility (Fig. 5B).

These results confirm that macrolide antibiotics exert a selective pressure both in pwCF and PCD leading to the accumulation of uL4 and uL22 mutations causing antibiotic resistance.

## Discussion

Antimicrobial resistance poses one of the greatest challenges to public health^31^. In the case of macrolides and *P. aeruginosa* infections, this is exacerbated by the lack of specific knowledge on the mode of action *in vivo* and on the unrecognized ability of infecting bacteria to develop antibiotic resistance ^14^. The assumption that *P. aeruginosa* is intrinsically resistant to macrolides underlines the limitations in mimicking antibiotic efficacy within the unique infectious microenvironment and modelling susceptibility accurately ^32,33^. This misconception - based on standardized AST - has inadvertently allowed the emergence of macrolide-resistant bacteria, which evade detection since routine AST for macrolides on *P. aeruginosa* is not commonly performed in clinical laboratories. Given their continuous use for the treatment of pwCF, COPD and PCD, it is imperative to recognize macrolide antibiotics’ effectiveness against *P. aeruginosa* and implement improved protocols for the early detection of resistant strains. We previously identified clinical strains of *P. aeruginosa* harbouring uL4 mutations resulting in significantly increased macrolide resistance ^14^. However, clinical strains harbour a multitude of SNPs and indels, which makes it challenging to disentangle the individual contributions of each mutation to the observed phenotype ^34,35^. Hence, isolating the effects of specific ribosomal mutations by comparing them in an isogenic background is crucial. This approach has allowed us to investigate the direct effects of representative uL4 and uL22 mutations in comparison to the parental wild-type strain on antibiotic susceptibility but also any collateral phenotypes. As hypothesized, the selected uL4 and uL22 mutations conferred increased resistance to erythromycin and azithromycin when moved into the PAO1 genetic background. Additionally, the macrolide-induced changes in phenotypes such as oxidative stress response and quorum sensing required a higher concentration of antibiotic in the mutant strains compared to the wild-type PAO1. Azithromycin efficacy on *P. aeruginosa* wild-type and two selected mutant strains was confirmed by performing bacterial infections in an ALI model system of pseudostratified airway epithelium in the presence of antibiotics ^15^. Azithromycin’s bacteriostatic effect was, indeed, considerably higher for the wild-type PAO1 strain than the ΔRA and +G strains. These results confirm that macrolides are effective antibiotics against *P. aeruginosa*’s infections. Moreover, the uL4 and uL22 ribosomal protein mutations not only serve to reduce the direct antibacterial effect of macrolide antibiotics but also mitigate the increase in redox sensitivity and the quorum sensing modulation, which is normally elicited by macrolide treatment.

Importantly, mutations in the ribosome can change the translational landscape resulting in alterations of growth characteristics and phenotypes ^1,3^. Some of the uL4 and uL22 mutant strains showed reduced growth rates which indicate a fitness cost associated with such mutations. Notably, in clinical strains of *P. aeruginosa* infecting pwCF, reduced growth rates are advantageous, facilitating persistence and evasion of host defences ^25,36,37^. Moreover, proteomic profiling revealed additional alterations in the expression of virulence factors such as those of the T3SS and T6SS which also contribute to the persistent phenotype ^37^. This reduced virulence was further confirmed in the ALI infection model with the uL4 mutant strains perturbing the epithelial cell layer to a lesser extent than their wild-type parental strain. Importantly, uL4-dependent changes in virulence profiles in clinical isolates of *P. aeruginosa* were not observed before ^14^. This is likely because years of adaptive evolution in patients might have drastically modified the phenotype and virulence profile of clinical strains before the accumulation of ribosomal mutations ^37^. Regulation of virulence in *P. aeruginosa* is complex and requires the integration of both intracellular and extracellular signals ^38,39^. Interestingly, the observed changes in the expression of virulence determinants in uL4 and uL22 mutants resemble the switch between acute and chronic infection phenotypes ^37^ regulated by the Gac/Rsm signaling pathway ^40^. However, we did not find changes in expression levels of components of this regulatory pathway. On the contrary, it has been shown in several bacterial species that mutations in ribosomal proteins or associated factors conferring antibiotic resistance to ribosome-targeting antibiotics can alter virulence ^41–44^. This suggests a generalized mechanism of virulence pathways expression dependent on the ribosome functionality and cell homeostasis possibly associated with the bacterial translation stress response activated as a consequence of the ribosomal mutations ^45^. Still, further analyses are required to determine the cross-regulation between ribosome mutations and virulence and this can only be revealed by the use of complex model systems taking into account the complexity of an infection and the host-pathogen interactions. ALI airway models can fulfil such role and allow more in-depth investigations of antibiotic resistance, virulence and persistence ^46,47^. For future advances, the integration of an active immune system will be beneficial for even closer to patient investigations.

The persistence of uL4 and uL22 mutant strains – as evidenced by isolation from the same patient over up to 8 years – confirms that any fitness costs, which the mutation may cause, do not lead the strains to be outcompeted. On the contrary, the combination of increased macrolide resistance, reduced virulence factor expression and slower growth could provide a competitive advantage allowing these strains to persist for years in the patient while also evading detection through reduced growth and because *P. aeruginosa* is typically not sampled with the intent of detecting macrolide resistance ^36^. It is, therefore, highly clinically relevant to inform and promote an accurate use of macrolides to avoid the emergence of new and highly persistent macrolide-resistant strains. In other words, it is not without cost to apply macrolide treatment even when the spectrum of activity is not believed to cover the pathogen in question.

In conclusion, our findings highlight the importance of systematic investigations of bacterial pathogens under *in vivo*-like conditions to comprehensively understand antibiotic efficacy, identify mechanisms of antibiotic resistance, and update antibiotic use guidelines which take into consideration secondary effects of antibiotic resistance mutations. These efforts will contribute to a new understanding of treatment failure and provide new therapies, which are essential for combatting the growing threat of increasing antibiotic resistance.

## Material and Methods

### Bacterial strains and growth conditions

The *P. aeruginosa* laboratory strain PAO1 was used as reference strain for the construction of recombinant strains ^48^. *P. aeruginosa* clinical isolates were isolated at the Department of Clinical Microbiology, at Rigshospitalet, Copenhagen, Denmark. Analyses of the bacterial isolates were approved by the local ethics committee of the Capital Region of Denmark (Region Hovedstaden; registration numbers H-1-2013-032, H-4-2015-FSP). Strains were routinely grown in LB medium at 37°C. For AST bacteria were grown in 50% LB medium (Sigma-Aldrich, St. Louis, Missouri, USA) (LB diluted with sterile water).

### Recombinant strains construction

Derivatives of pACRISPR were constructed with the Uracil Specific Excision Reagent (USER) cloning ^49^. The plasmids used in this study are listed in Table S3, and the primers in Table 4. Briefly, target DNA fragments were amplified from the respective clinical strain genome with the corresponding primers using Phusion U polymerase kit (Thermo Fisher Scientific, USA), cloned into the pACRISPR plasmid using the USER® Enzyme (New England Biolabs, USA) and transformed into *Escherichia coli* DH5alpha cells. PCR was performed with the random colonies selected in LB agar plates with 50 µg/ml carbenicillin using OneTaq® 2X Master Mix (New England Biolabs, USA) and sequenced with Sanger method (Eurofins Scientific, Luxembourg) to confirm the correct insertions. *P. aeruginosa* genome editing was carried out according to the following protocol ^50^. Briefly, cells were transformed with pCasPA plasmid via electroporation, as previously described ^51^ and selected on LB agar plates with 50 µg/ml tetracycline. The resulting strains were further transformed with the appropriate derivative of pACRISPR plasmid (Table S3) using a similar method but with the addition of 0.2% arabinose in the growth medium, and transformants were selected in plates with 50 µg/ml tetracycline and 150 µg/ml carbenicillin. For the second crossover, the resulting colonies were grown in liquid LB at 37°C for 2h, afterwards arabinose was added to 0.2% to induce the expression of the editing system. Cells were further incubated for 2h and streaked on an LB agar plate containing 5% sucrose. On the next day, colony PCR following Sanger sequencing Eurofins Scientific, Luxembourg) was used to confirm the presence of desired mutations.

### Susceptibility testing

MICs were determined by standard broth microdilution ^52^ with minor modifications ^14^. Briefly, overnight (o.n.) cultures grown in LB broth were diluted to a density of approximately 5×10^5^ cells/ml in 50% LB broth. 190 μl bacterial cell culture was dispensed into a 96-well plate along with 10 μl antibiotic stock solution and incubated in a BioTek ELx808 plate reader with shaking at 37°C for 24 h, where optical density (OD) was measured every 20 min at 630 nm. The MIC was determined as the lowest concentration, which inhibited visible growth (OD_630nm_ < 0.05) in the wells. After incubation, 5 μl of each well was spotted onto LB-agar plates to determine the post-MIC effect defined as the lowest concentration of antibiotic that inhibits regrowth on plates after o.n. incubation.

E-tests (bioMérieux, Marcy-l’Étoile, France) were performed according to producer’s guidelines but on 50% LB agar instead of standard cation-adjusted Müller-Hinton broth. Briefly, colonies were picked from fresh LB-plates and resuspended into PBS for a density of 10^8^ CFU/ml. Sterile cotton swabs were used to spread the suspension evenly onto 50% LB plates, which were allowed to dry for 5-10 min before application of the E-test. Results were read after 24h incubation at 37°C. Growth curves from the untreated cultures from the MIC determination were used for the determination of the generation time. Maximum generation time was computed directly after the end of the lag-phase by fitting an exponential growth equation to 3-7 OD datapoints using GraphPad Prism 8 version 8.4.3.

### Swimming assay

Swim agar was prepared by mixing 50% LB and 0.3% agar. 25 ml plates were poured with or without azithromycin in the concentration of 0.25% the MIC of the mutant strains. After drying for 1h at room temperature, plates were point inoculated with the test strain and incubated top-up at 30°C for 24h after which the swimming diameter was measured. For each strain, three independent biological replicates were analyzed.

### Twitching motility

Strains were stab inoculated onto the bottom of a thin 1.5% LB agar plate. After 24h incubation at 37°C in a humidified box, the LB agar was carefully removed, and the twitch zone visualized by staining with 0.1% crystal violet solution followed by 2 washes with mQ water.

### Biofilm formation

O.n. cultures were diluted to an OD_600nm_ of 0.1 in 50% LB and diluted 10^3^, 10^4^ or 10^5^-fold before 150 µl was aliquoted into a Calgary Biofilm Device type plate and incubated statically for 24h at 37°C. Bacterial growth was measured in the plate at OD_630nm_. The PEG-lid was washed 3x in 175 µl mQ water before staining the biofilm with 200 µl 0.1% crystal violet solution for 30 min. After 3 washes in 200 µl mQ water, the lid was dried for 30 min, destained in 30% acetic acid, and the wells measured for absorbance at 600nm to quantify the crystal violet. Each measurement was normalized to the bacterial growth.

### Redox sensitivity

Strains were grown for 24h at 37°C in 5 ml 50% LB medium containing 0% or 2% of the MIC concentration of azithromycin. 100 μl of o.n. cultures (corresponding to OD_600nm_ = 1) was mixed with 3 ml molten 0.5% agar, 50% LB and poured onto pre-cast LB-agar plates. Over-lay agar was set to solidify for 5-10 min at room temperature whereafter a disc of filter paper saturated with 5 μl fresh 30% H_2_O_2_ was placed on top. After 24h at 37°C, clearing zones around the H_2_O_2_ discs were measured. For each strain, three independent biological replicates were analyzed.

### Protease activity and pyocyanine secretion

Strains were grown for 24h in 5 ml 50% LB medium containing 0% or 2% of the MIC concentration of azithromycin. 1 ml of o.n. culture was harvested, and the supernatant filtered through a 0.22 µm filter. The cell-free sterile supernatant was collected and 2 μl spotted onto 1% skimmed milk plates in 1% agar. Clearing zones were measured after 24h at 37°C. The same supernatant was used for pyocyanine semi-quantification. Absorbance at 690nm was measured on 200 μl of the filtered supernatant in a BioTEK Synergy multimode reader.

### Pyoverdine production

O.n. cultures were diluted and treated as described for the broth microdilution 96-well plates MIC assay either without antibiotic or including 2% of the azithromycin MIC of mutant strains. Pyoverdine production was measured in the 96-well plates at 405nm excitation and 460nm emission after 20h incubation at 37°C in a BioTEK Synergy multimode reader and normalized to the OD at 630nm.

### Proteomic analysis

Frozen cell pellets were thawed on ice and any remaining supernatant was removed after centrifugation at 15,000g for 10 min. While kept on ice, two 3-mm zirconium oxide beads (Glen Mills, NJ, USA) were added to the samples. Immediately after moving the samples away from ice 100 μl of 95 °C GuanidiniumHCl (6M Guanidinium hydrochloride (GuHCl), 5mM tris(2-carboxyethyl)phosphine (TCEP), 10 mM chloroacetamide (CAA), 100 mM Tris–HCl pH 8.5) was added to the samples. Cells were disrupted in a Mixer Mill (MM 400 Retsch, Haan, Germany) set at 25 Hz for 5 min at room temperature, followed by 10 min in thermo mixer at 95° at 2000 rpm. Any remaining cell debris was removed by centrifugation at 15,000g for 10 min, after which 50 μl of supernatant were collected and diluted with 50 μl of 50 mM ammonium bicarbonate. Based on protein concentration measurements (BSA), 100 μg of protein was used for tryptic digestion. Tryptic digestion was carried out at constant shaking (400) for 8h, after which 10 μl of 10% TFA was added and samples were ready for StageTipping using C18 as resin (Empore, 3M, USA). For analysis, a CapLC system (Thermo scientific) coupled to a Orbitrap Q-exactive HF-X mass spectrometer (Thermo Scientific) was used. First samples were captured at a flow of 10 μl/min on a precolumn (µ-precolumn C18 PepMap 100, 5 µm, 100Å) and then at a flow of 1.2 µl/min the peptides were separated on a 15 cm C18 easy spray column (PepMap RSLC C18 2 µm, 100Å, 150 µmx15 cm). The applied gradient went from 4% acetonitrile in water to 76% over a total of 60 min. MS-level scans were performed with Orbitrap resolution set to 60,000; AGC Target 3.0e^6^; maximum injection time 50 ms; intensity threshold 5.0e^3^; dynamic exclusion 25 sec. Data dependent MS2 selection was performed in Top 20 Speed mode with HCD collision energy set to 28% (AGC target 1.0e4, maximum injection time 22 ms, Isolation window 1.2 m/z). After acquisition, the raw data were analysed using the Proteome discoverer 2.4 software with the following settings: Fixed modifications: Carbamidomethyl (C)I and Variable modifications: oxidation of methionine residues. First search mass tolerance 20 ppm and a MS/MS tolerance of 20 ppm. Trypsin as enzyme and allowing one missed cleavage. FDR was set at 0.1%. The Match between runs window was set to 0.7 min. Quantification was only based on unique peptides and normalization between samples was based on total peptide amount. For the searches the Pseudomonas aeruginosa PAO1 reference proteome (UniProt Proteome ID UP000002438) was used. Protein abundances were log_2_ transformed and differentially expressed proteins were analysed by two-way ANOVA with Tukey’s multiple comparisons test. Proteins with a Log_2_(Fold-Change) ≥ |0.6| and Adjusted P value ≤ 0.05 were considered differentially expressed. Principal Component Analysis (PCA), and hierarchical clustering analysis were performed using the software JMP Pro version 17. Enrichment analysis was performed using the DAVID Functional Annotation Bioinformatics Microarray Analysis tool from lists of Locus Tags of proteins that were differentially expressed.

### Air Liquid Interface infection model preparation

ALI infection models were established using BCi-NS1.1 cells ^53^, as previously outlined ^15^. Initially, cells were expanded in Pneumacult-Ex Plus medium (STEMCELL Technologies) within culture flasks in a humidified incubator at 37°C with 5% CO_2_. Following expansion, 10^5^ cells were seeded onto 1 μm pore polyester membrane inserts (Falcon) coated with type I collagen (Gibco), and incubated until reaching full confluency in a humidified incubator at 37°C with 5% CO_2_. Subsequently, ALI conditions were initiated by aspirating the media from the apical chamber and replacing the basolateral chamber media with Pneumacult-ALI maintenance medium (STEMCELL Technologies) supplemented with 480 ng/mL hydrocortisone, 4 µg/mL heparin (STEMCELL Technologies), Pneumacult-ALI maintenance supplement (STEMCELL Technologies) and Pneumacult-ALI 10x supplement (STEMCELL Technologies). Differentiation of ALI cultures was achieved by incubating the cells at 37°C with 5% CO_2_ in a humidified incubator for 28 days, with media replacement every 3-4 days. Epithelial polarization was assessed by regular measurements of TEER using a chopstick electrode (STX2; World Precision Instruments). Following the initial 15 days under ALI conditions, the apical surface was washed with PBS every 7 days to remove accumulated mucus.

### Infection procedure and characterization

O.n. cultures of *P. aeruginosa* were employed to inoculate 4 mL of LB medium at an OD_600nm_ of 0.01, and grown until reaching mid-exponential phase. Subsequently, 1 mL of the culture was centrifuged at 5000 rpm for 5 minutes, and the bacterial pellets were washed with PBS and resuspended in PBS to a concentration of 10^5^ CFUs/mL. A 10 μL aliquot of this suspension, containing 10^3^ CFUs, was inoculated onto the apical surface of fully differentiated BCi-NS1.1 ALI cultures, while the basolateral chamber contained 600 µL of supplemented Pneumacult-ALI maintenance medium. Control wells received an equal volume of bacteria-free PBS. The infected cells were then incubated at 37 °C with 5% CO_2_ in a humidified incubator for 14 h, after which 200 µL of PBS were added to the apical surface and transepithelial electrical resistance TEER was measured.

The 200 µL of PBS from the apical surface and the 600 µL of basolateral medium were collected for determination of CFUs by plating 10 µL of serial dilutions on LB-agar plates. For quantification of bacterial CFUs attached to the epithelium, 200 μL of PBS were added to the apical surface, and the epithelium was disrupted using a cell scraper. CFUs in the resulting solution were quantified by plating on LB-agar plates. The basolateral media harvested at the end of the infection was also used for measuring LDH by using the Invitrogen™ CyQUANT™ LDH Cytotoxicity Assay Kit (Invitrogen), in technical triplicates, following the manufactureŕs instructions.

For confocal microscopy examination of the infection process, infected ALI cultures were rinsed with PBS and fixed with 4% paraformaldehyde in both apical and basolateral chambers for 20 min at 4 °C. Following fixation, cells were washed with PBS and permeabilized and blocked with a buffer containing 3% BSA, 1% saponin, and 1% Triton X-100 in PBS for 1 hour. Staining was performed by incubating cells for 2 h at room temperature with a solution of Phalloidin-AF555 (Invitrogen) and TO-PRO3 (Biolegend) diluted 1:400 and 1:1000, respectively, in staining buffer (3% BSA and 1% Saponin in PBS). Fixed and stained epithelia containing transwells were removed from their supports with a scalpel and mounted on glass slides with VECTASHIELD® Antifade Mounting Medium. Imaging of the epithelia was performed using a Leica Stellaris 8 Confocal Microscope (40× magnification, 1.3 oil) and images were analyzed using the LasX software (Leica).

To assess the efficacy of azithromycin against bacteria colonizing the epithelium in the infection model, first, the infection was initiated as described previously. After 7 h of incubation at 37°C with 5% CO_2_ in a humidified incubator, azithromycin was introduced into the basolateral medium to a final concentration of 512 µg/mL. The system was then further incubated for 14 h in presence of azithromycin. Subsequently, TEER was measured, and the amount of CFUs in the apical side, basolateral medium, or attached to the epithelium was determined by plating 10 µL of serial dilutions from the corresponding harvested bacterial suspension onto LB-agar plates, following the procedure described earlier. Confocal microscopy analysis of the infection process under azithromycin treatment was conducted by processing the samples as described above.

### Whole Genome Sequencing of uL4 and uL22 recombinant strains

The genome of the recombinant strains was sequenced to confirm the presence of the specific ribosome mutation. gDNA of the recombinant *P. aeruginosa* strains was extracted using the DNeasy Blood & Tissue Kit (Qiagen). Library preparation and sequencing were performed by Novogene (UK) Company Limited. Briefly, gDNA was sheared into short fragments. The obtained fragments were end-repaired, A-tailed and further ligated with Illumina adapter. The fragments with adapters were PCR amplified, size selected, and purified. The library was checked with Qubit and real-time PCR for quantification and bioanalyzer for size distribution detection. Sequencing reads were trimmed, and low-quality reads and potential contamination from adapters were removed using Trimmomatic (v 0.35) tool ^54^. Genomic analysis was conducted by BacDist ^55^ to identify genomic variants relative to PAO1 reference genome (NCBI: NC_002516.2). BacDist filtered mutations to only retain variants with a mapping quality of at least 50, a minimum coverage of 10, and a minimum fraction of 50% of reads supporting the variant at a given position.

### Identification of uL4 and uL22 mutations

The sequences of the genes encoding uL4 (*rplD*) and uL22 (*rplV*) were extracted from a dataset comprising 663 full genome sequenced *P. aeruginosa* genomes, which are available upon request. This dataset included 431 isolates obtained from pwCF and 232 isolates obtained from PCD patients who were treated at Rigshospitalet, Copenhagen, Denmark. Nucleotide sequences were extracted from these genomes and aligned using the MUSIAL tool ^56^. Mutations were subsequently identified by comparing these sequences to the reference strain PAO1.

### Data availability

The authors declare that all data necessary for supporting the findings of this study are enclosed in this paper (Supplementary Data 1 and Supplementary Table 1). Whole genome sequencing data of the recombinant strains is publicly available through the SRA number XXX. Sequencing data on the 663 clinical isolates is available upon request from the corresponding author.

## Acknowledgements

We thank the Infection Microbiology group for their insightful comments and discussion. We also thank Victor Mora for performing the preliminary ALI infections.

## Author’s contribution

LG, RLR, HKJ and SM designed the project. OI performed the whole genome sequencing data analysis, RLR performed the proteomics data analysis, PL, LG and IP performed the experiments. HKJ provided strains, clinical data and funding. SM provided funding. LG and RLR verified the data and wrote the first draft of the paper. All authors approved the final version of the manuscript.

